# Anesthetic-like effects of ketamine in *C. elegans*

**DOI:** 10.1101/2024.08.28.610063

**Authors:** Katariina Seppälä, Inés Reigada, Olli Matilainen, Tomi Rantamäki, Leena Hanski

## Abstract

Transparency of *C. elegans* enables microscopic *in vivo* imaging of cellular processes, but immobilization is required due to high locomotor activity. Preservative NaN_3_ is commonly used for this, but is associated with oxidative stress and toxicity. Here, anesthetic-like effects of dissociate anesthetic ketamine in *C. elegans* are presented using video recordings and infrared-based automated activity tracking. Ketamine caused a reversible blockade of locomotion at a similar concentration (20-50 mM) at which NaN_3_ produces paralysis. Moreover, the recovery rate was remarkably high, and short-term ketamine treatment did not show signs of SKN-1 activation, a marker of the stress response.

## 1 INTRODUCTION

The nematode *Caenorhabditis elegans* expresses homologue proteins of the key mammalian targets of general anesthetics^1^, including gamma-aminobutyric acid type A receptors (GABA_A_-R)^2^ and glutamatergic *N*-methyl-D-aspartate receptors (NMDAR)^3^, making it a relevant model organism to study mechanisms of general anesthetics^4,5^. Due to *C. elegans*’ susceptibility to anesthesia, these agents could be used for imaging, as they can induce loss of locomotion in the worm^4^. Thus far, research has focused on GABAergic gaseous anesthetics^4,5^, but extending research to include other types of anesthetics could enhance this application. Methods for inducing immobility in *C. elegans* include immobilizing chemicals, agarose pads, microfluidic devices and cold shock^6^. NaN_3_, a commonly used chemical agent for *C. elegans* immobilization, is highly toxic, which inevitably activates stress response pathways mediated by SKN-1^6^. SKN-1, an orthologue of the mammalian Nrf/CNC proteins, is a crucial transcription factor in *C. elegans*, regulating detoxification and metabolic functions in oxidative and other stress responses^7^. Consequently, the use of NaN_3_ may confound certain experimental results.

In this study, the potential of ketamine, an NMDAR blocking anesthetic, to chemically immobilize *C. elegans* was investigated.

## 2 MATERIALS AND METHODS

### 2.1 Nematode cultures

Previously described methods were applied for the maintenance and handling of *C. elegans*^8^. Briefly, N2 wild type Bristol and CL2166 dvIs19 [(pAF15)gst-4p::gfp::nls] (Caenorhabditis Genetics Center (CGC), University of Minnesota, Minneapolis, MN, USA) strains were maintained at 17 °C on Nematode Growth Medium (NGM; USbiological Life sciences, Swampscott, MA, USA) agar plates seeded with *E. coli* OP50 (CGC). Egg-lay synchronized young adults were used for all experiments. Sterile S Basal buffer (100 mM NaCl, 6 mM K_2_HPO_4_, 40 mM KH_2_PO_4_; pH 7) was used for washing and as assay medium and negative control in all assays, unless otherwise stated.

### 2.2 Pharmaceutical agents

Ketamine-HCl solution (Ketaminol Vet 50 mg/mL, Intervet International B.V. Boxmeer, Netherlands), NaN_3_ powder (Acros Organics Bvba, Geel, Belgium) and 30% H_2_O_2_ solution (Sigma-Aldrich, St Saint Louis, MO, USA) were used.

### 2.3 Preliminary immobilization assays

Immobilization of the worm was used as a quantitative endpoint, defined as a lack of full body movement for 10 seconds^9^. Head tossing was not regarded as mobility. NaN_3_ at concentration of 20 mM was used as a positive control^6^. The worms were observed in room temperature under Leica S9D imaging system (Leica Microsystems, Wetzlar, Germany) and manually scored as mobile or immobile.

### 2.4 Video recording of ketamine-induced anesthetic-like responses

N2 worms were individually picked with a platinum worm pick from a maintenance plate into 96-well plates containing different concentrations of ketamine (1-50 mM), 20 mM NaN_3_ and control. Each worm was recorded with Evos M7000 Cell Imaging System (Thermo Fisher Scientific, Waltham, MA, US) and manually scored in one of 3 categories: immobile, uncoordinated or thrashing. The videos were analyzed blinded to avoid researcher bias.

### 2.5 Automatized activity measurement

The motility of *C. elegans* N2 was studied using wMicroTracker ONE (Phylumtech, Santa Fe, Argentina). This system measures the locomotory activity of worms by detecting interruptions to infrared beams in multiwell plates containing worms in liquid media. Worms were individually picked from maintenance plates into 96-well plates containing test solutions, and their activity was recorded for 60 minutes at intervals of 15 minutes.

### 2.6 Recovery assays

N2 worms were exposed to the ketamine treatments, 20 mM NaN_3_ and S-basal negative control in a 96-well plate for 30 minutes. Subsequently, the worms were transferred onto *E. coli* NGM plates and manually rated as mobile or immobile at indicated time points. The worms’ survival was recorded 24 hours later. Worms were counted as dead if they did not move spontaneously or when prodded with a platinum worm pick.

### 2.7 Radial dispersion assay

Radial dispersion assay was modified from previously described method for gaseous anaesthetics^9^. N2 worms were washed from maintenance plates and incubated in ketamine solutions for 30 minutes at room temperature. Volumes containing approximately 50 worms were transferred onto NGM plates seeded with a 2 mm ring of *E. coli* OP50 overnight preculture along the outer edge as chemoattractant. The plates were incubated for 45 minutes at room temperature. The percentages of worms having reached the peripheral ring of *E. coli* were counted manually under a microscope.

### 2.8 SKN-1 activation

CL2166 worms were washed from maintenance plates and treated with test solutions for 30 minutes. S-basal buffer and 10 mM H_2_O_2_ were used as negative and positive controls, respectively and subjected for the GFP readout as previously described^10^. The worms were transferred to Eppendorf tubes and frozen for protein extraction as described^11^, using Pierce BCA Protein Assay Kit (#23227, Thermo Fisher). Fluorescence values were normalized with the protein concentrations.

### 2.9 Statistical analysis

GraphPad Prism was used for data handling and statistical analysis using unpaired Student’s t-test or Mann-Whitney U-test, when applicable.

## 3 RESULTS

### 3.1 Ketamine-induced immobility in *C. elegans*

Effects of ketamine on N2 worm mobility maintained in liquid media in 96-well plates were initially screened by visual inspection at 0.5-50 mM concentrations range 30-minutes post-incubation. Majority of the worms showed no locomotion at 20 mM or higher concentrations. At these concentrations a subset of worms displayed sluggish, uncoordinated movements, but no spatial movement. Concentrations ≤5 mM did not cause observable changes in worm motility.

Next, we conducted video recordings to analyse the effects of ketamine. The behaviour of the worms was rated as immobile, uncoordinated (displaying irregular movement) or “thrashing” (i.e. fast, lateral swimming movement^12^). The worms lost their characteristic thrashing movement at concentrations of ≥5 mM ketamine (**Figure 1**). Notably, while NaN_3_ yielded full paralysis, ketamine-induced changes in the nematode motor functions affected mostly locomotion. At all concentrations, there was individual variation in response to ketamine, but immobility and sluggish movement increased dose-dependently. The ketamine-induced dose-dependent decrease in locomotor activity was also quantitatively assessed with wMicrotracker ONE infrared microbeam detection system. Here, the worm motility was significantly affected already at concentrations ≥5 mM. At 20 mM, the average motility scores approached total inactivity in this setup, illustrating the lack of locomotion. The levels of immobilization at 20 mM and 50 mM enabled fluorescent and brightfield imaging with the fluorescent strain CL2166 (**Figure 1E**).

**Figure 1.**
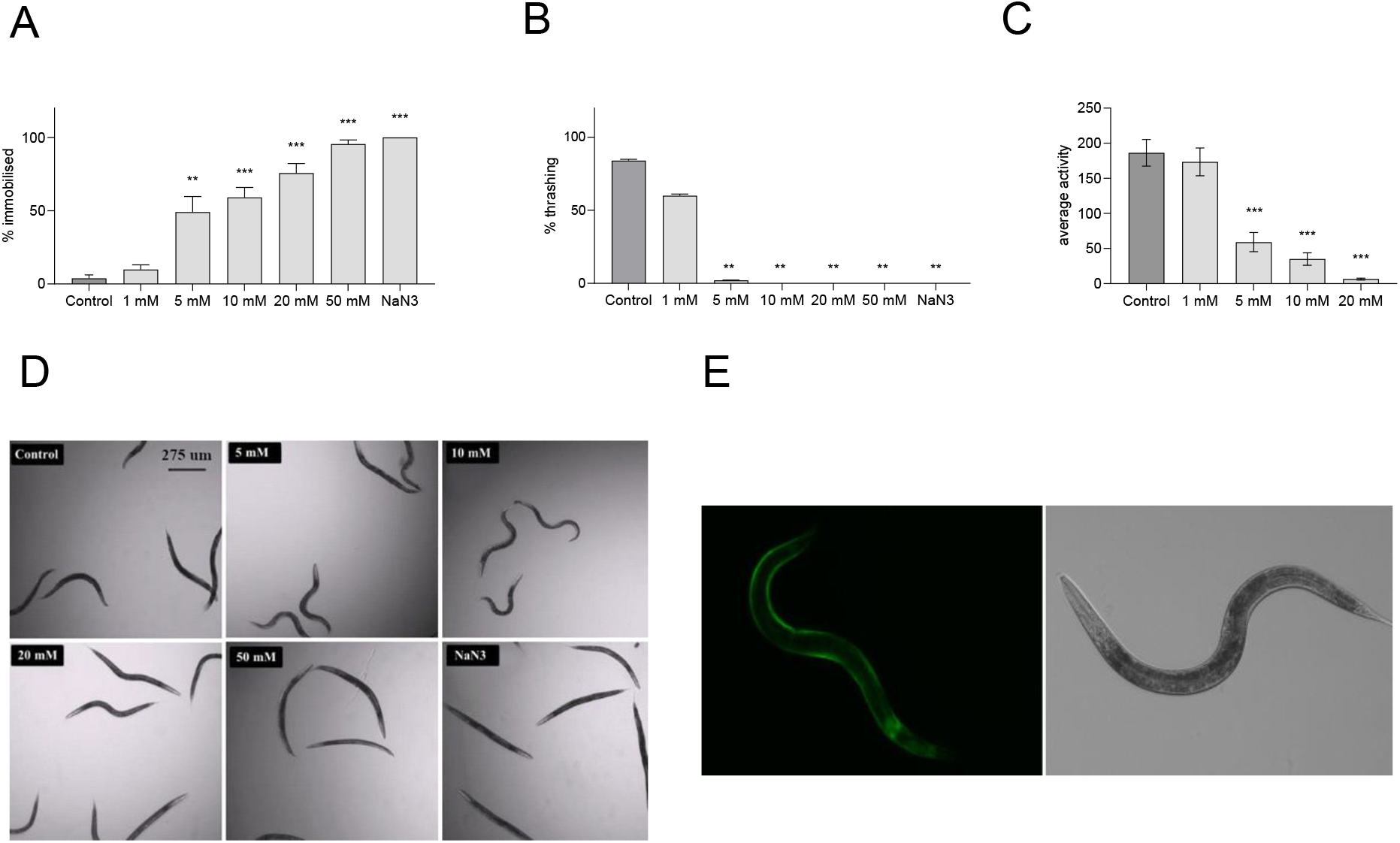
The impact of ketamine on the locomotion of *C. elegans*. A) Immobilization of *C. elegans* N2 treated with different concentrations of ketamine in 96-well plates. B) Thrashing locomotion of *C. elegans* N2 treated with different concentrations of ketamine in 96-well plates. C) wMicrotracker ONE measured 30-minute locomotor activity scores of *C. elegans* N2 treated with different concentrations of ketamine in 96-well plates. D) Images of *C. elegans* N2 treated with negative control, different concentrations of ketamine and 20 mM sodium azide in a 96-well plate. Magnification: 4X. E) Fluorescent and brightfield images of *C. elegans* CL2166 immobilized with 20 mM ketamine. Magnification: 10X. Figures show pooled data from 2 individual experiments with a minimum of 2 technical replicates per experiment, n ≥ 50 for all test groups (n = total number of worms). Data are presented as mean ± SEM. Asterisks (***: *p*<0.001, **: *p*<0.01, *: *p*<0.05) indicate statistically significant differences with the untreated control in unpaired Student’s t test (A, C) and Mann-Whitney U test (B).

### 3.2 Recovery, survival, and oxidative stress in response to ketamine

The recovery of mobility in *C. elegans* N2 at different time points post-treatment is seen in **Figure 2**. Most worms treated with 20 mM ketamine recovered within 30 minutes, resuming their characteristic, undulating crawling locomotion. The speed of recovery was comparable to that of the worms treated with 20 mM NaN_3_. Recovery from 50 mM ketamine was altered even at 3 hours post-exposure, without influencing short-term (24 h) mortality. However, radial dispersion assay, which measures worms’ ability to find a peripheral food source^4^, revealed marked chemosensation deficiency after 20 mM (but not lower) ketamine exposure. This indicates prolonged disruption of chemosensation at this dose, despite normalization of mobility.

**Figure 2.**
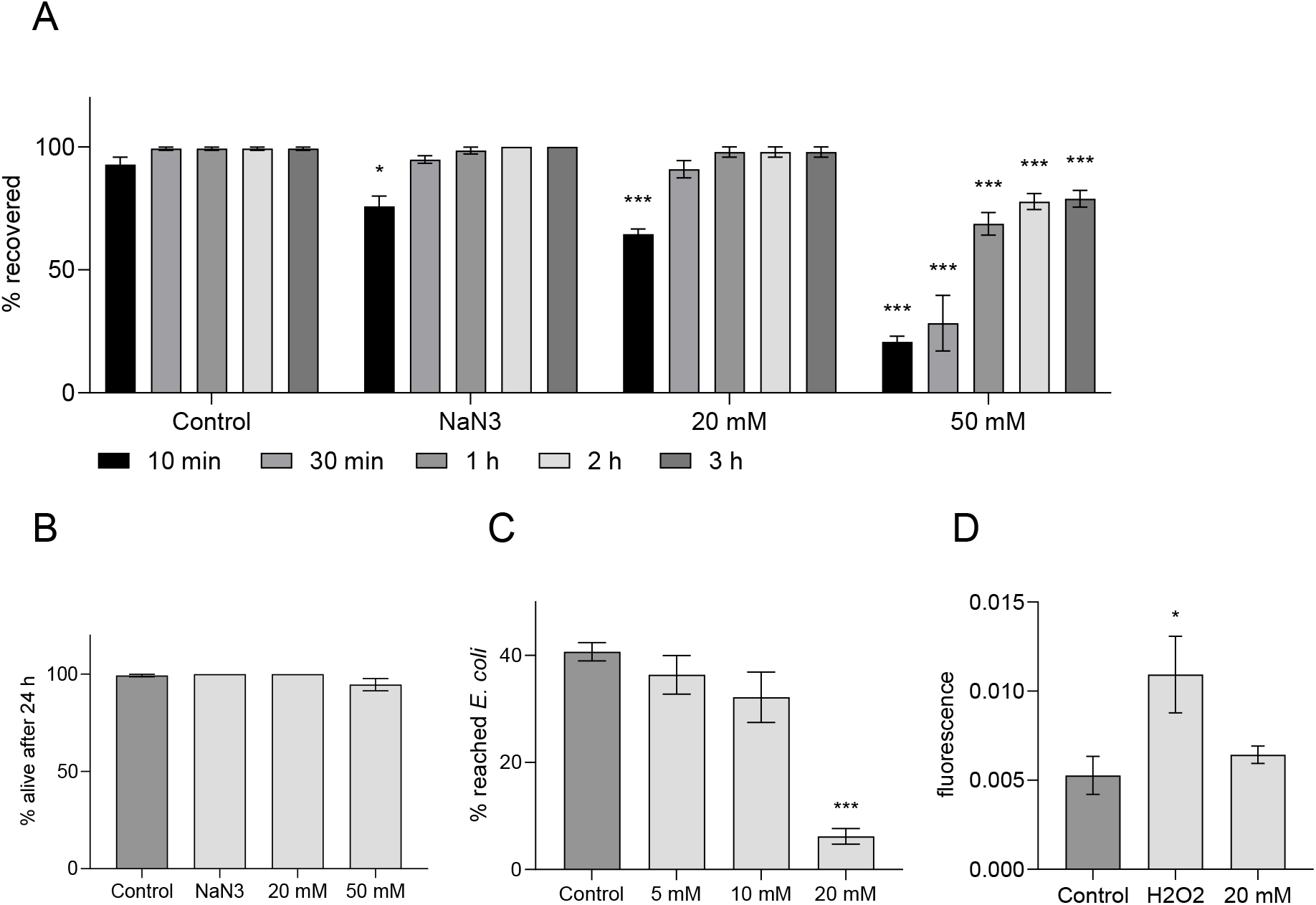
Recovery post ketamine treatment and the effects of ketamine on radial dispersion and SKN-1 mediated stress response in *C. elegans*. A) *C. elegans* N2 recovery from immobilization by different concentrations of ketamine, n ≥ 56 for all groups (n = total number of worms). B) Survival of *C. elegans* N2 24 hours after treatment by different concentrations of ketamine, n ≥ 54 for all groups. C) Chemoattraction to *E. coli* of *C. elegans* N2 treated with different concentrations of ketamine, n ≥ 200 for all groups. D) Protein normalised fluorescence values of *C. elegans* CL2166 treated with negative control, positive control (H_2_O_2_) and 20 mM ketamine. Figures represent pooled data from 2 individual experiments with a minimum of 2 technical replicates. Data are shown as mean ± SEM. Asterisks (***: *p*<0.001, **: *p*<0.01, *: *p*<0.05) indicate statistically significant differences to the control treatment (unpaired Student’s t test).

Ketamine-induced changes in SKN-1 activation were studied in CL2166 mutants that carry a transcription reporter construct of GFP (green fluorescent protein) to the GST-4 (glutathione S-transferase) promoter and show increased GFP levels when transcription factor SKN-1 is activated^10^. The values of the ketamine (20 mM) treated group were not significantly higher compared to the negative control, while H_2_O_2_ produced significant increase.

## 4 DISCUSSION

Our results suggest that ketamine has potential as a non-toxic immobilizing agent for *C. elegans*. NaN_3_, an inhibitor of mitochondrial respiration, induces hypoxia in *C. elegans*^6^. It is toxic and has been shown to cause pathological changes in *C. elegans* tissues, such as elevated expression of stress proteins and activation of SKN-1 mediated signaling^6^. Unlike NaN_3_, ketamine does not produce full paralysis but still induces a level of immobilization compatible with imaging without marked SKN-1 activation.

The findings correspond to the literature on *C. elegans* locomotory responses to GABAergic volatile anesthetics^5^. Further studies are however required to elucidate underlying mechanisms of ketamine in *C. elegans*. The anesthetic mechanism of action of ketamine is linked mainly to NMDAR antagonism, but other mechanisms such as inhibition of nicotinic and muscarinic acetylcholine receptors may also be involved^13^. In *C. elegans*, NMDAR subunit homologues NMR-1 and NMR-2 are expressed predominantly in command interneurons regulating forward and backward movement of the worm^3^. NMDAR homologues are also associated with memory retention in learned avoidance responses^14^. Nitrous oxide (N_2_O), another NMDAR antagonist, requires NMR-1 for its action in *C. elegans*^15^, but induces only behavioral defects, not immobility, in *C. elegans*. Loss-of-function mutations of the NMDAR subunit homologues cause defects in learning, forward-backward movement and foraging behavior, but they do not severely disrupt development or locomotion^14^.

In conclusion, ketamine appears to be a non-toxic alternative for immobilizing *C. elegans*, offering immobilization for imaging without triggering significant stress responses.

## Acknowledgements

This study has been supported by Academy of Finland (grant no. 332920 to O.M. and 333291 and 358425 to L.H.) and Sigrid Juselius Foundation (T.R.).

## Authorship contribution statement

**Katariina Seppälä**: Writing – review & editing, Writing – original draft, Visualization,, Methodology, Investigation, Formal analysis, Conceptualization, Data curation. **Inés Reigada:** Writing – review & editing, Methodology, Investigation, Formal analysis, Data curation. **Olli Matilainen:** Methodology, Review & editing, Funding acquisition. **Tomi Rantamäki:** Writing – review & editing, Supervision, Funding acquisition, Project administration. **Leena Hanski:** Writing – review & editing, Supervision, Funding acquisition, Project administration.

